# Evaluation of common *in vitro* assays for the prediction of oral bioavailability and hepatic metabolic clearance in humans

**DOI:** 10.1101/2024.02.25.581937

**Authors:** Urban Fagerholm

## Abstract

**Introduction:** Intrinsic hepatic metabolic clearance (CL_int_) measured with human hepatocytes, apparent intestinal permeability (P_app_) obtained using the Caco-2 model, unbound fraction in plasma (f_u_) and blood-to-plasma concentration ratio (C_bl_/C_pl_) are commonly used for predicting the hepatic clearance (CL_H_) and oral bioavailability (F) of drug candidates in humans. The primary objective was to select drugs whose *in vitro* hepatocyte CL_int_, Caco-2 P_app_, f_u_ and C_bl_/C_pl_ have been measured in various laboratories and studies, and estimate correlation coefficients (R^2^) for predicted and observed F and log plasma CL_H_. Secondary aims were to estimate the laboratory/study variability and its impact on predictions and to compare results to *in silico* and animal model-based predictions.

**Materials and Methods:** A literature search was done in order to find unbound hepatocyte CL_int_, (and corresponding predicted *in vivo* CL_int_), Caco-2 P_app_, f_u_ and C_bl_/C_pl_ data. Compounds with multiple measurements for the four assays, without significant *in vivo* solubility/dissolution limitations and with known *in vivo* CL_H_ and F, were selected. Min, max and mean estimates were used in the analysis.

**Results and Discussion:** Thirty-two compounds with data (in total 561 estimates) produced by 21 major pharmaceutical companies and universities met the inclusion criteria. The predicted vs observed R^2^ for log mean CL_int_, log mean CL_H_ and mean F were 0.32, 0.08 and 0.20, respectively. Exclusion of atenolol increased the R^2^ for CL_H_ to 0.20. R^2^-values were considerably lower than those presented in many studies, which seems to be explained by selection bias (choosing favorable reference values). There was considerable interstudy variability for measured and predicted CL_int_ (80- and 1,476-fold mean and max differences, respectively) and measured f_u_ (6.6- and 50-fold mean and max differences, respectively). For F, higher predictive performance was found for *in silico* (Q^2^=0.58; head-to-head) and animal *in vivo* models (R^2^=0.30).

**Conclusion:** The combination of data from many laboratories and the use of mean values resulted in reduced selection bias and predictive accuracy. Overall, the predictive accuracy (here R^2^) for log CL_int_, log CL_H_ and F was low to moderately low (0.08-0.32). The halved R^2^ compared to individual studies where high performance was demonstrated seems to be explained be selection bias (enabled by large data variability). Animal *in vivo* models, and in particular, *in silico* methodology, outperformed *in vitro* methodology for the prediction of F in man.

## Introduction

Intrinsic hepatic metabolic clearance (CL_int_) measured with human hepatocytes, apparent intestinal permeability (P_app_) measured using the Caco-2 cell model, unbound fraction in plasma (f_u_) and blood-to-plasma concentration ratio (C_bl_/C_pl_) can be used for predicting the oral bioavailability (F) of drug candidates in humans. Such studies and reports are, however, rare. It is more common that human hepatocyte CL_int_, f_u_ and C_bl_/C_pl_ are used for prediction of the hepatic metabolic clearance (CL_H_).

Some reports in the literature show high correlation coefficients (R^2^) (up to 0.7-0.9; with slopes close to 1) between predicted and observed log *in vivo* CL_int_ and CL_H_ (Riley et al. 2005; Hallifax et al. 2010; Sohlenius-Sternbeck et al. 2010; Yamagata et al. 2017), whereas other show low to moderate correlations (down to ca 0.2; with slopes clearly deviating from 1) (Stringer et al. 2008; Wood et al. 2017).

Variability within and between laboratories and differences between *in vitro* and *in vivo* conditions are known sources behind uncertainties and errors of *in vivo* predictions. For example, 3- to 3.5-fold mean interlaboratory variability (calculated as ratios between maximum and minimum values for each compound) was found for human hepatocyte CL_int_, Caco-2 P_app_ and f_u_ (Fagerholm 2022). An extensive investigation of interlaboratory variability of human hepatocyte CL_int_ and unbound CL_int_ by Louisse et al. 2020 showed coefficients of variance between 26 and 231 % and that values for the majority of compounds differed by more than one order of magnitude.

*In vitro* to *in vivo* predictions of *in vivo* CL_int,_ CL_H_ and F are sensitive to subjective bias (for example, selection of favorable or unfavorable f_u_- and C_bl_/C_pl_-data). Our previous investigation showed the impact of selecting favorable data for maximizing the shown predictive accuracy of *in vivo* CL_int_ (Fagerholm et al. 2022). It was possible to select f_u_- and C_bl_/C_pl_-data to reach a R^2^ for CL_int_ of ca 0.7-0.9, but also to reach R^2^s below 0.1. With the use of mean f_u_- and C_bl_/C_pl_-data a R^2^ of 0.38 was reached for CL_int_ (n=17).

The primary objective of the present study was to find and select drugs whose *in vitro* hepatocyte CL_int_, (and corresponding predicted *in vivo* CL_int_), Caco-2 P_app_, f_u_ and C_bl_/C_pl_ have been measured in various laboratories and studies, and estimate the R^2^ for predicted vs observed F and plasma CL_H_. Secondary aims were to estimate the laboratory/study variability and its impact on predictions and to compare results to *in silico* and animal model-based predictions.

## Materials & Methods

The literature was searched for studies with data for human hepatocyte CL_int_-predicted *in vivo* unbound CL_int_, Caco-2 P_app_ and *in vivo* fraction absorbed (f_a_), f_u_, C_bl_/C_pl_, non-renal CL (surrogate for CL_H_), F and gut-wall bioavailability (F_gut_).

Suitable data were found in the following references – Predicted unbound *in vivo* CL_int_ (Albaugh et al, 2012; Bowman and Benet 2019; Keefer et al. 2023), Caco-2 P_app_ (Skolnik et al. 2010; Lin et al. 2011, McGinnity et al. 2007, Thomas et al. 2005, Matsson et al. 2005, Lee et al. 2017), Li et al. 2018, Gertz et al. 2010, Sköld et al. 2006, Irvine et al. 1999, Yee 1997, Lau et al. 2004, Fujikawa et al. 2005, Li et al. 2007), f_u_ (Fagerholm et al. 2021a; Wang et al. 2014; Hallifax et al. 2010; Soars et al. 2002; Sohlenius-Sternbeck et al. 2010; Obach 1999; Obach et al. 2008; Yau et al. 2017; Yamagata et al. 2017; Keefer et al. 2023, Prosilico AB databank), C_bl_/C_pl_ (Hallifax et al. 2010; Sohlenius-Sternbeck et al. 2010; Obach 1999; Yamagata et al. 2017; Keefer et al. 2023, Yau et al. 2017; Uchimura et al. 2010; Prosilico AB databank), CL_H_ (Hallifax et al. 2010; Sohlenius-Sternbeck et al. 2010; Obach 1999; Yamagata et al. 2017; Keefer et al. 2023; Obach et al. 2008; Prosilico AB databank), F (Varma et al. 2010; Yau et al. 2017; Musther et al. 2014; Fagerholm et al. 2021b; Prosilico AB databank), and F_gut_ (Yau et al. 2017).

Caco-2 P_app_ *vs in vivo* f_a_-relationships were established for each report with Caco-2 P_app_-data (including data for many other reference substances), and f_a_ for the selected compounds was predicted based in P_app_-f_a_-relationships obtained for each study.

Only compounds with complete *in vivo* dissolution and *in vitro* CL_int_>limit of quantification (LOQ) were selected.

Selected data were produced by companies including Astellas Research Institute, AstraZeneca, Boehringer-Ingelheim, Chugai, Cyprotex, GSK, Millennium Pharmaceuticals, Novartis, Pfizer, Roche and Schering Plough, RIKEN Innovation Center, and Universities of Chungnam, Kyoto, Manchester, New Jersey, Nottingham, Shanghai, South Dakota, Sungkyunkwan and Uppsala.

Plasma CL_H_ and the corresponding F_H_ were estimated using the well-stirred liver model with a liver blood flow rate of 1500 mL/min and by considering C_bl_/C_pl_ for F-predictions. The F was estimated as f_a_ • F_gut_ • F_H_, where f_a_ was predicted from P_app_ *vs* f_a_-relationships in each Caco-2 study.

For each selected compound, minimum, maximum and mean F- and plasma CL_H_-values were estimated, and predicted mean values were compared to corresponding measured *in vivo* mean values. To further evaluate the results for compounds with known gut-wall metabolism by CYP3A4 and available *in vivo* F_gut_-estimates these were either removed or their F_gut_ was considered.

*In silico* (forward-looking results by Prosilico AB; head-to-head) and animal model (mice, rats and dogs) predicted data were taken from Fagerholm et al. 2021 and compared to *in vitro* data based predictions.

## Results & Discussion

### Selected compounds and variability between studies and data sets

Thirty-two compounds that met the inclusion criteria were found and selected. Their measured and predicted data are shown in Tables 1 and 2. In total, 175 predicted *in vivo* CL_int_, 126 P_app_ (over 500 estimates including references), 172 and 88 f_u_ and C_bl_/C_pl_-values were found and used (in total 561 estimates), which corresponds to an average of 5.5, 3.9, 5.4 and 2.8 CL_int_-, P_app_-, f_u_- and C_bl_/C_pl_-estimates per compound, respectively.

**Table 1.**
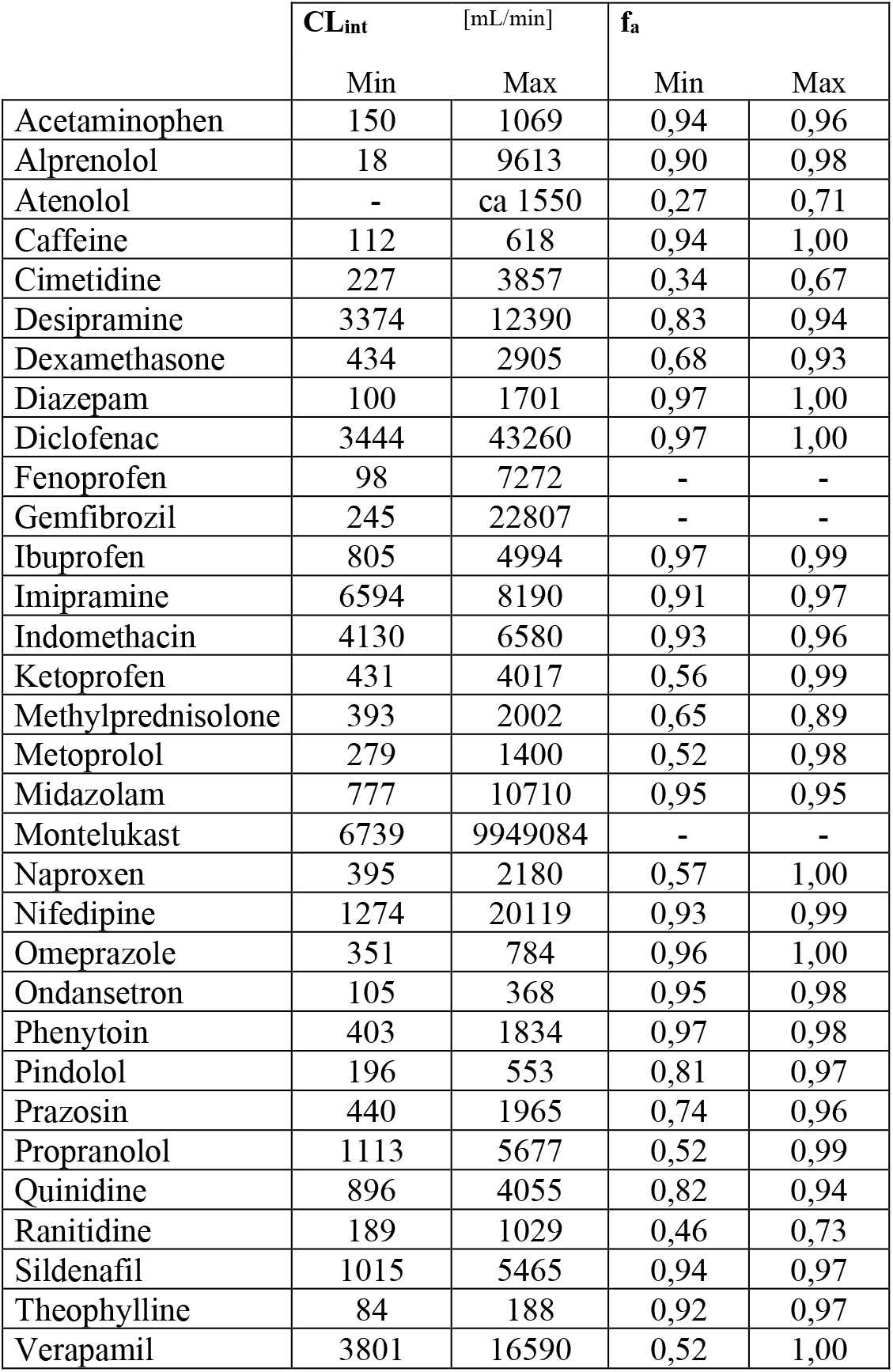
The selected compounds and their CL_int_ (predicted from human hepatocyte unbound CL_int_) and f_a_ (predicted from Caco-2 P_app_).

**Table 2.** The selected compounds and their predicted and observed plasma CL_H_ and F and measured F_gut_.

|  | Plasma CL<sub>H</sub> [mL/min] |  |  |  | F |  |  |  | F<sub>gut</sub> |
| --- | --- | --- | --- | --- | --- | --- | --- | --- | --- |
|  | Predicted |  |  | Observed | Predicted |  |  | Observed | Observed |
|  | Min | Max | Mean | Mean | Min | Max | Mean | Mean |  |
| Acetaminophen | 71 | 545 | 206 | 340 | 0,60 | 0,92 | 0,83 | 0,82 |  |
| Alprenolol | 3 | 803 | 411 | 1008 | 0,27 | 0,98 | 0,63 | 0,08 | CYP3A4 |
| Atenolol | - | 757 | 645 | 2,1 | 0,14 | 0,71 | 0,32 | 0,50 |  |
| Caffeine | 68 | 372 | 167 | 97 | 0,65 | 0,95 | 0,85 | 0,98 | CYP3A4 |
| Cimetidine | 109 | 1013 | 506 | 14 | 0,10 | 0,62 | 0,35 | 0,68 | CYP3A4 |
| Desipramine | 259 | 897 | 605 | 756 | 0,32 | 0,84 | 0,61 | 0,40 |  |
| Dexamethasone | 54 | 574 | 300 | 225 | 0,35 | 0,89 | 0,65 | 0,80 | CYP3A4 |
| Diazepam | 1 | 59 | 12 | 27 | 0,90 | 1,00 | 0,97 | 0,93 | CYP3A4 |
| Diclofenac | 6,86 | 1114 | 664 | 243 | 0 | 0,99 | 0,26 | 0,54 |  |
| Fenoprofen | 0,65 | 133 | 44 | 118 | - | - | - | - |  |
| Gemfibrozil | 0,49 | 531 | 191 | 217 | - | - | - | - |  |
| Ibuprofen | 5 | 86 | 35 | 57 | 0,87 | 0,99 | 0,94 | 0,98 |  |
| Imipramine | 321 | 722 | 512 | 892 | 0,45 | 0,84 | 0,66 | 0,53 |  |
| Indomethacin | 16 | 457 | 191 | 77 | 0,42 | 0,95 | 0,79 | 0,93 |  |
| Ketoprofen | 2 | 208 | 58 | 78 | 0,42 | 0,98 | 0,85 | 0,85 |  |
| Methylprednisolone | 54 | 352 | 217 | 406 | 0,49 | 0,86 | 0,70 | 0,90 | CYP3A4 |
| Metoprolol | 113 | 684 | 462 | 865 | 0,30 | 0,92 | 0,66 | 0,40 |  |
| Midazolam | 13 | 509 | 112 | 371 | 0,37 | 0,94 | 0,84 | 0,36 | 0,57 |
| Montelukast | 6,04 | 1478 | 1421 | 48 | - | - | - | - |  |
| Naproxen | 0 | 38 | 11 | 5 | 0,54 | 1,00 | 0,91 | 0,86 |  |
| Nifedipine | 10 | 716 | 189 | 511 | 0,16 | 0,98 | 0,76 | 0,50 | 0,68 |
| Omeprazole | 14 | 51 | 31 | 582 | 0,91 | 0,99 | 0,92 | 0,54 | CYP3A4 |
| Ondansetron | 28 | 214 | 116 | 386 | 0,79 | 0,96 | 0,88 | 0,60 | CYP3A4 |
| Phenytoin | 20 | 197 | 153 | 33 | 0,76 | 0,96 | 0,82 | 0,85 |  |
| Pindolol | 106 | 355 | 259 | 266 | 0,54 | 0,91 | 0,73 | 0,87 |  |
| Prazosin | 13 | 150 | 44 | 316 | 0,62 | 0,95 | 0,84 | 0,68 |  |
| Propranolol | 78 | 764 | 373 | 756 | 0,20 | 0,94 | 0,66 | 0,24 | CYP3A4 |
| Quinidine | 96 | 575 | 254 | 238 | 0,42 | 0,87 | 0,70 | 0,73 | 0,90 |
| Ranitidine | 132 | 592 | 279 | 154 | 0,27 | 0,67 | 0,50 | 0,60 |  |
| Sildenafil | 22 | 291 | 102 | 637 | 0,65 | 0,95 | 0,86 | 0,41 | 0,72 |
| Theophylline | 33 | 106 | 61 | 54 | 0,84 | 0,94 | 0,90 | 0,90 | CYP3A4 |
| Verapamil | 231 | 1060 | 703 | 1222 | 0,04 | 0,85 | 0,41 | 0,24 | 0,57 |

Estimate ranges for CL_int_, f_a_, f_u_, C_bl_/C_pl_, CL_H_ and F were 6-57,500 mL/min, 0.50-1.00, 0.0009-0.97, 0.56-1.67, 2-1222 mL/min and 0.08-0.98, respectively. Thus, there were wide estimate ranges for all parameters except for f_a_.

The average, median and maximum CL_int_-ratios (based on differences between studies) were 80-, 6.7- and 1,476-fold, respectively. Corresponding values for f_a_ were 1.34-, 1.13- and 2.63-fold, respectively. The average, median and maximum f_u_-ratios were 6.64-, 2.62- and 50-fold, respectively. Corresponding values for C_bl_/C_pl_ were 1.24-, 1.16- and 1.78-fold, respectively. Thus, there was considerable interstudy variability for measured and predicted CL_int_ and measured f_u_, which is in line with findings in previous reports (Bowman and Benet 2019; Fagerholm et al. 2021a; Fagerholm 2022).

More than 3-fold differences between maximum and minimum predicted F were found for 8 compounds – alprenolol (3.6-fold; predicted F=0.27-0.98; observed F=0.08), atenolol (5.0-fold; predicted F=0.14-0.71; observed F=0.50), cimetidine (6.5-fold; predicted F=0.10-0.62; observed F=0.68), diclofenac (>99-fold; predicted F=0-0.99; observed F=0.54), metoprolol (3.1-fold; predicted F=0.30-0.92; observed F=0.40), nifedipine (6.2-fold; predicted F=0.16-0.98; observed F=0.50), propranolol (4.7-fold; predicted F=0.20-0.94; observed F=0.24), verapamil (22-fold; predicted F=0.04-0.858; observed F=0.24).

Very large differences between maximum and minimum predicted CL_H_ were found for some compounds. For example, diclofenac (CL_H_ predicted to be low to high; 162-fold difference between lowest and highest predicted CL_H_), montelukast (CL_H_ predicted to be low to very high; 245-fold difference between lowest and highest predicted CL_H_), gemfibrozil (CL_H_ predicted to be very low to moderate; 1,083-fold difference between lowest and highest predicted CL_H_), fenoprofen (CL_H_ predicted to be very low to low; 205-fold difference between lowest and highest predicted CL_H_) and naproxen (CL_H_ predicted to be very low to low; 135-fold difference between lowest and highest predicted CL_H_).

The R^2^ for mean predicted *vs* observed f_a_ and log CL_int_ were 0.75 (Figure 1) (average R^2^=0.5 for all studies) and 0.32 (Figure 2), respectively. The moderately low predictive accuracy for hepatocyte-based predictions of *in vivo* log CL_int_ (R^2^=0.32) was similar to that found for 16 Pfizer drug discovery compounds by Keefer et al. 2023 (0.43) and by Stringer et al. 2008, but about half of that found by, for example, Sohlenius-Sternbeck et al. 2010 (0.72; 0.38 after correction for reference data) and Hallifax et al. 2010 (0.62).

**Figure 1.**
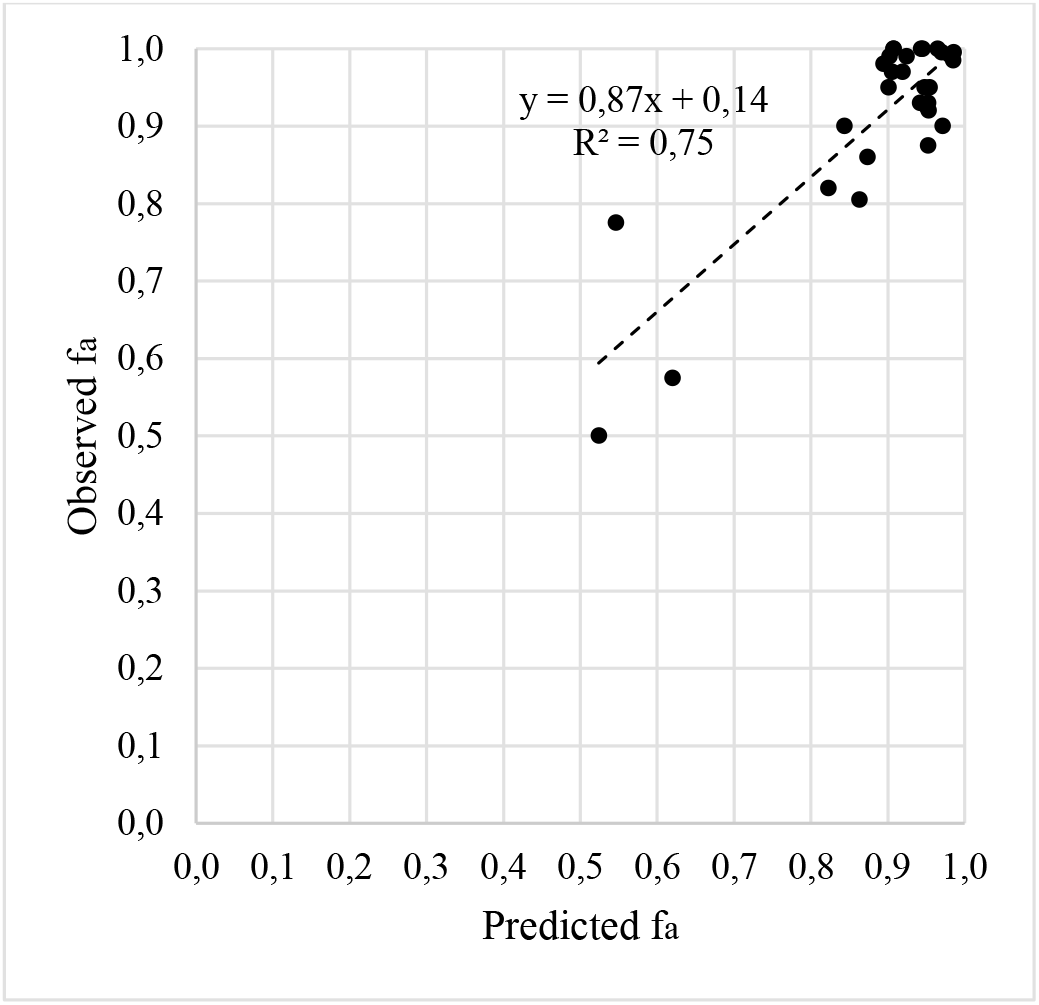
The correlation between predicted and observed mean f_a_ (n=29).

**Figure 2.**
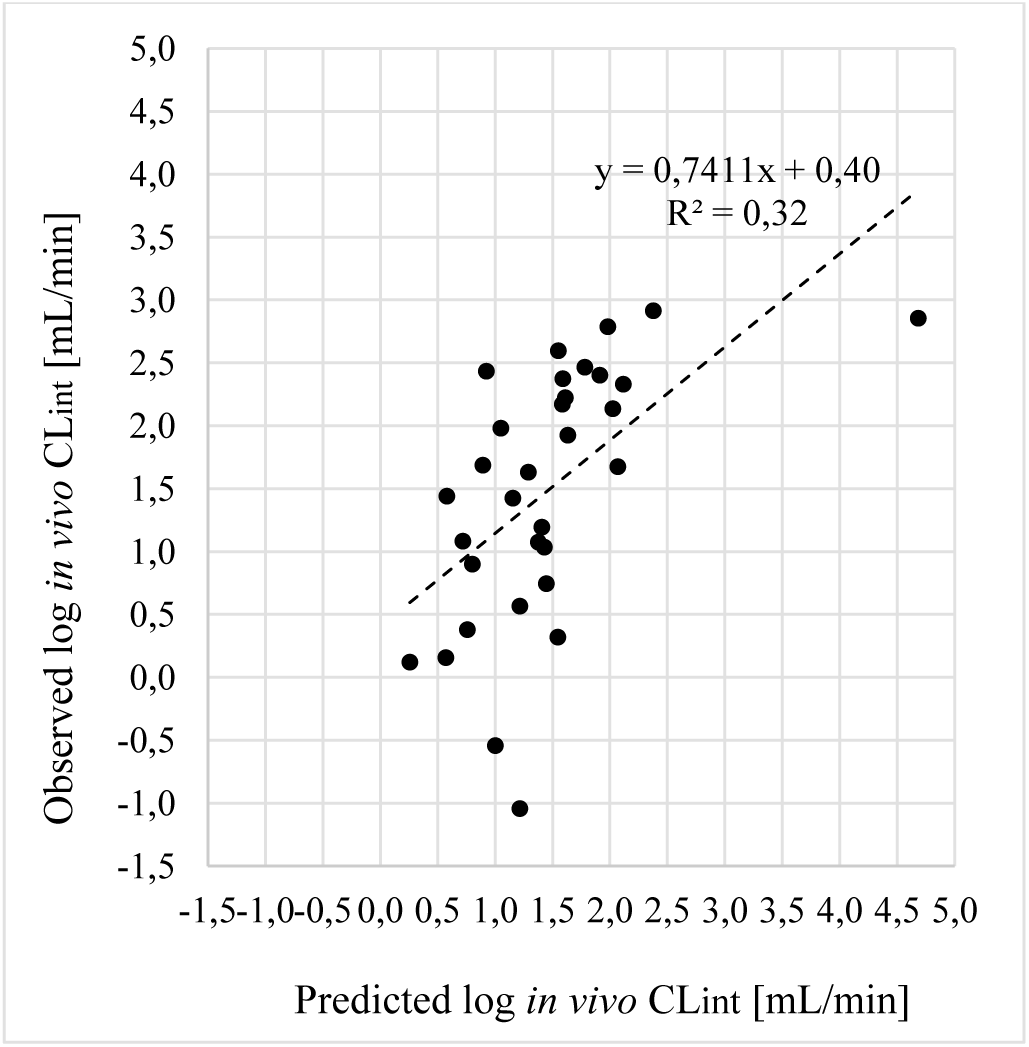
The correlation between predicted and observed mean log CL_int_ (n=32).

Based on the results it appears that a R^2^ of ca 0.3±0.2 is appropriate for human hepatocyte-based predictions of *in vivo* log CL_int_.

### Prediction of oral bioavailability

Twenty-nine of the 32 compounds qualified for evaluation of prediction of F. The R^2^ between predicted and observed mean F was 0.20 (Figure 3). Consideration of F_gut_ for and removal of 5 CYP3A4-substrates with known measure F_gut_ had only slight impact on the F-predictions (R^2^=0.26). Median and max ratios for highest and lowest predicted F were 2.1- and >99-fold, respectively. 72 % of observed F-estimates were within prediction ranges (Figure 4).

**Figure 3.**
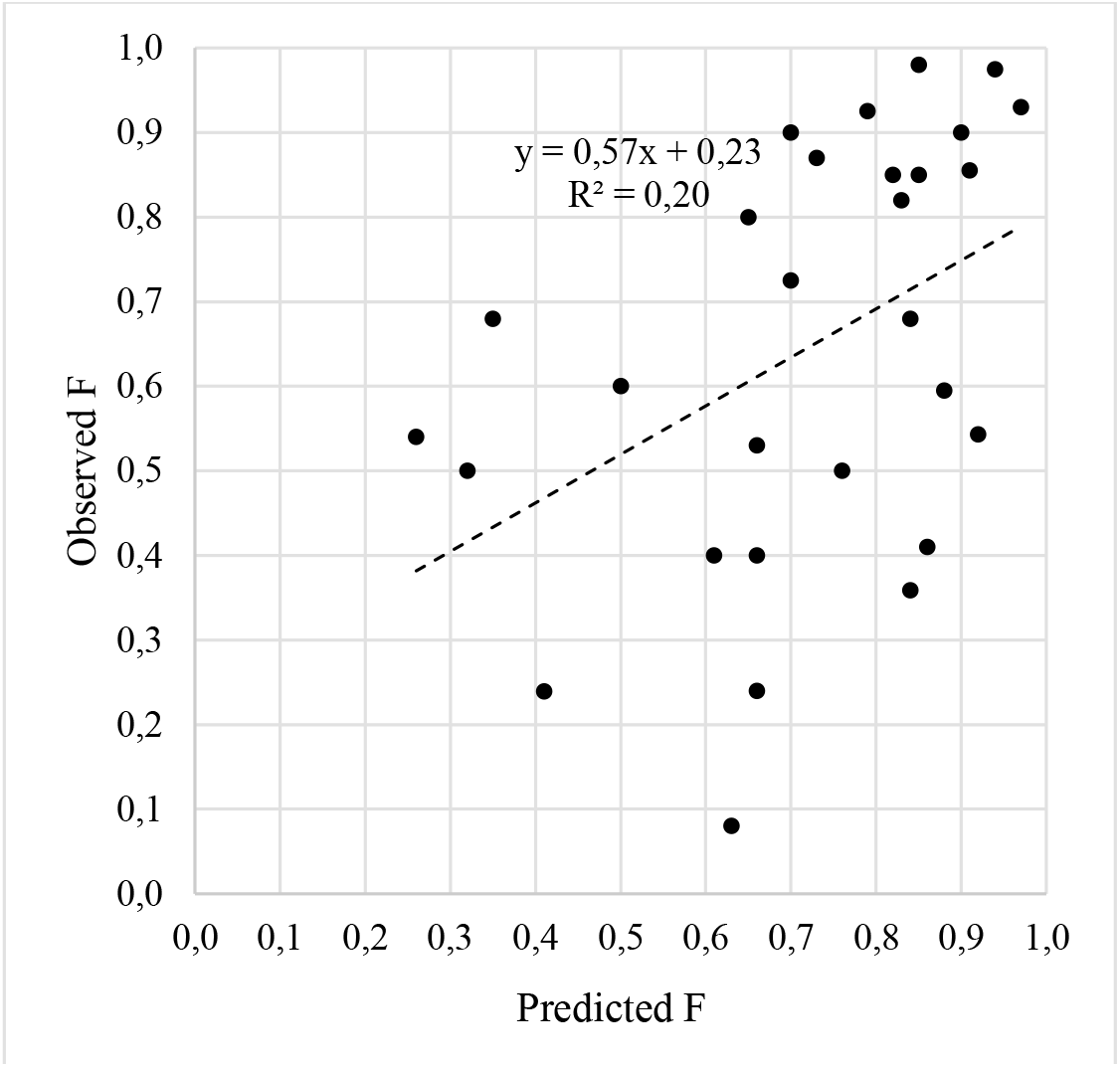
The correlation between predicted and observed mean F (n=29).

**Figure 4.**
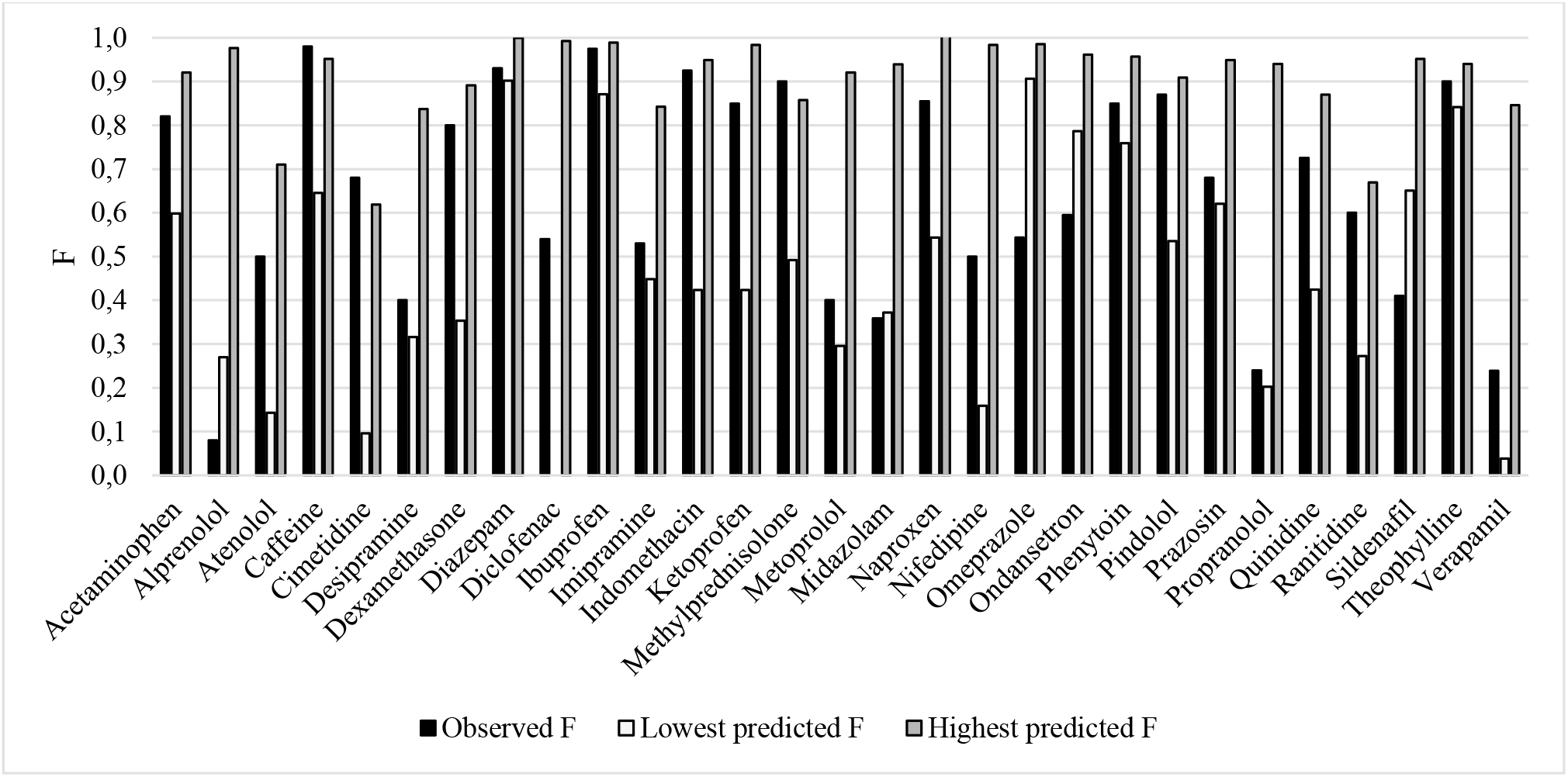
Observed mean F and predicted min and max F (n=29).

The R^2^ for F based on mean data from many sources was considerably lower than that reported from one laboratory by Li et al. 2007. With human hepatocyte CL_int_ and Caco-2 P_app_ they reached a R^2^ of 0.66 for 30 compounds (including 10 of the compounds in the present study).

### Prediction of plasma hepatic clearance

The R^2^ between predicted and observed mean log plasma CL_H_ was 0.08 (Figure 5). Exclusion of atenolol (with only one literature *in vitro* CL_int_-estimate) the increased the R^2^ for plasma CL_H_ to 0.20. Mean, median and max ratios for highest and lowest predicted CL_H_ were 81-, 10- and 1,083-fold, respectively. 69 % of observed CL_H_-estimates were within prediction ranges (Figure 6).

**Figure 5.**
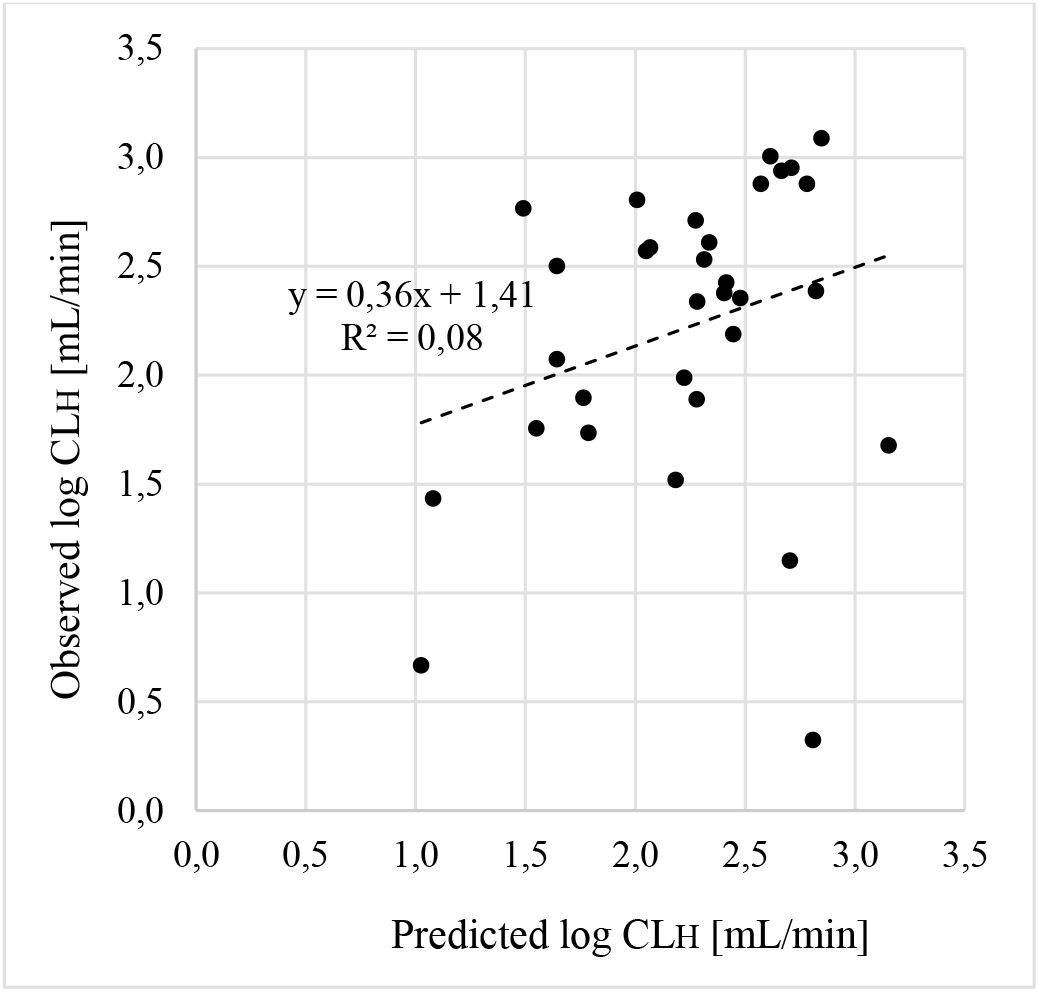
The correlation between predicted and observed mean log plasma CL_H_ (n=32).

**Figure 6.**
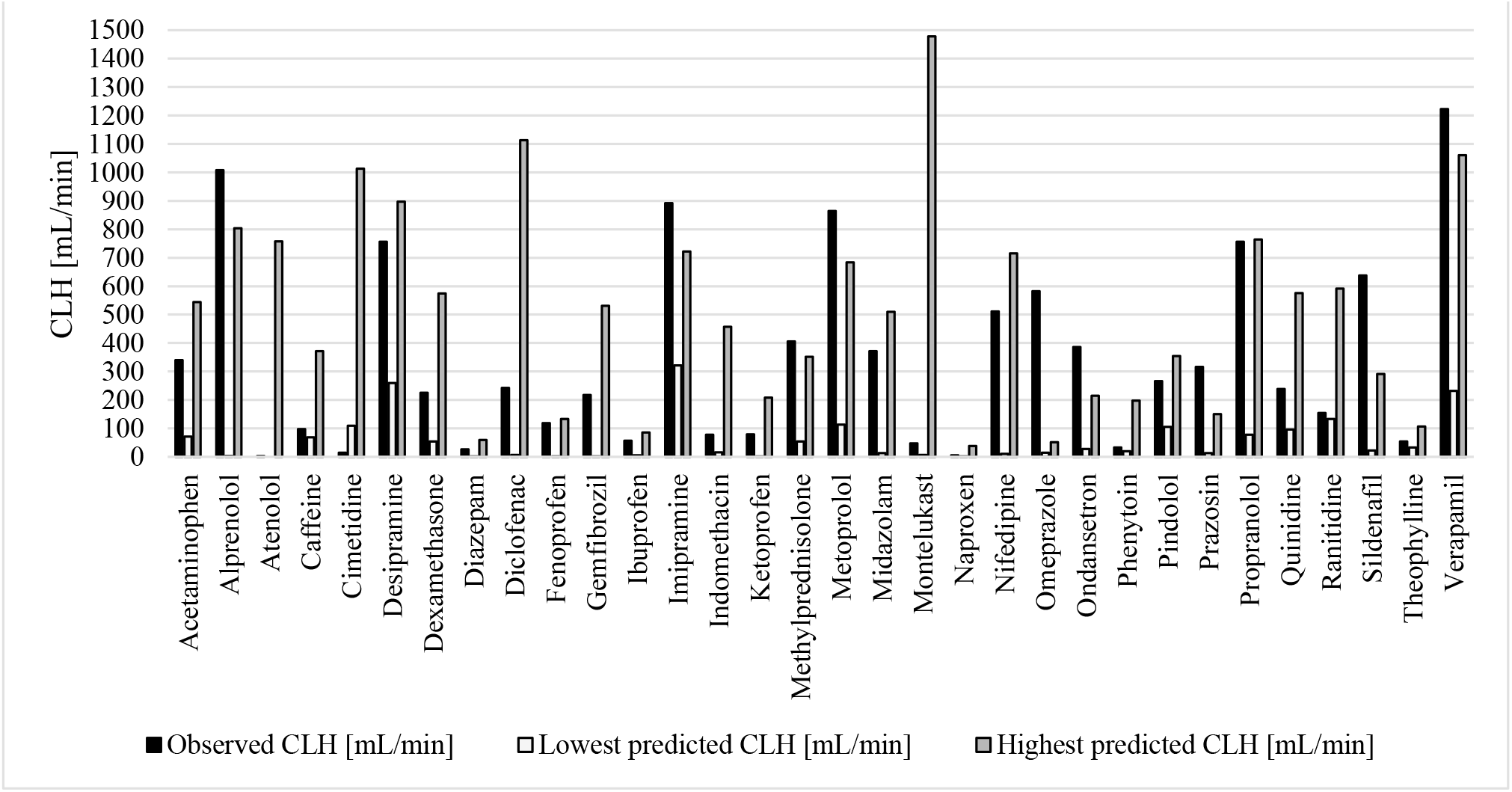
Observed plasma CL_H_ and predicted min and max plasma CL_H_ (n=32).

In line with what was found for F, the R^2^ for log plasma CL_H_ based on mean data from many sources was considerably lower than that reported by, for example, Riley et al. 2005; Hallifax et al. 2010, Sohlenius-Sternbeck et al. 2010 and Yamagata et al. 2017) (ca 0.7-0.9). It was closer to estimates by Stringer et al. 2008; Wood et al. 2017 and Fagerholm et al. 2021a and for a test set by Keefer et al. 2023 (ca 0.2-0.4). The low reproducibility and high predictive accuracy in some studies is interesting.

### The impact on data selection for predictions

The reason to higher R^2^-values in separate reports compared to those reached with mean data from multiple sources could be retrospective selection of favorable f_u_, C_bl_/C_pl_ and *in vivo* CL_H_-data. Support for that is given by interstudy differences in selected data by companies and researchers.

F_u_- and C_bl_/C_pl_-data for 23 compounds from 3 studies were used in prediction studies by one player in the pharmaceutical industry, and in 82 % of cases different data were used for each study (Figure 7). The mean and maximum interstudy differences for f_u_ were 1.89- and 4.50-fold, respectively. Corresponding estimates for C_bl_/C_pl_ were 1.14- and 1.59-fold, respectively. For every other compound it was possible to select f_u_- and C_bl_/C_pl_-data to adjust for 2- to 5-fold differences. Replacement of selected f_u_ and C_bl_/C_pl_ with average values resulted in large decrease of R^2^ and increase of prediction errors.

**Figure 7.**
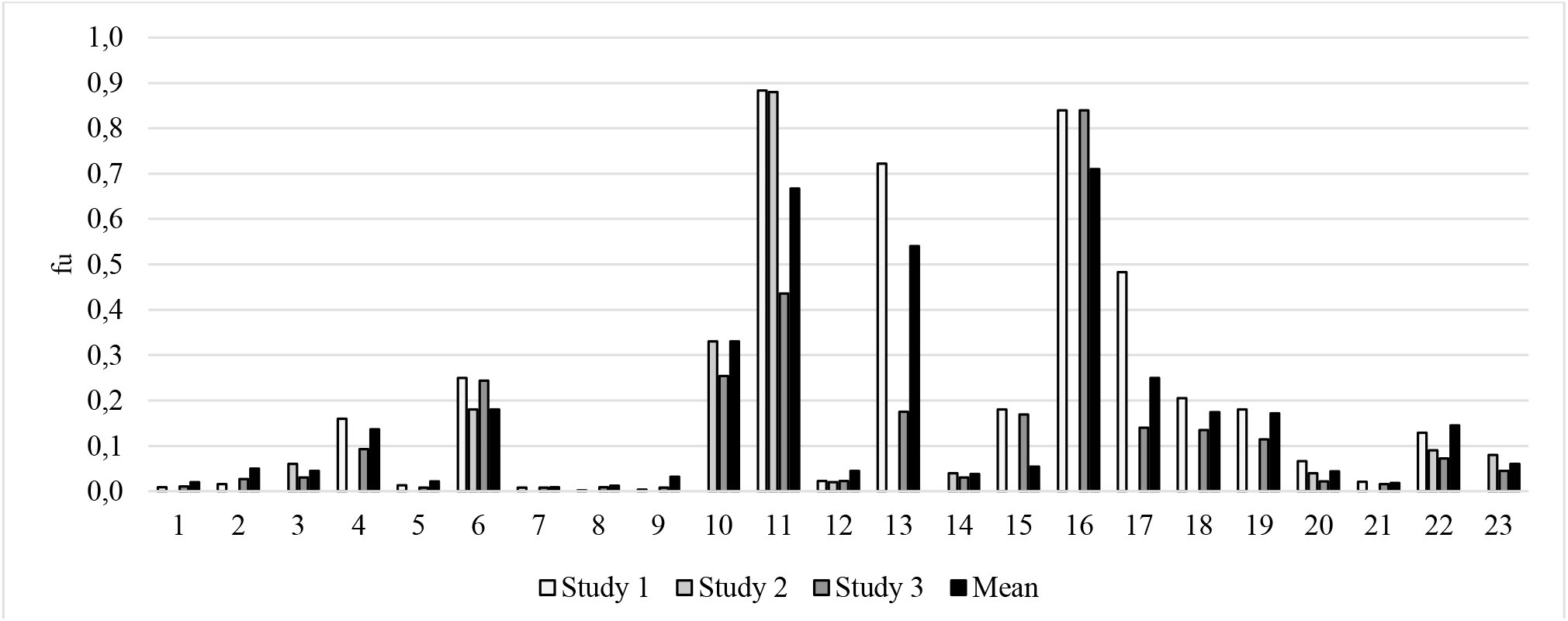
F_u_-values for 23 compounds selected in different studies by one player in the pharmaceutical industry, including mean f_u_-values estimated from more studies.

F_u_-data for 28 compounds from 4 studies were used in prediction studies by another player, and in all (100 %) cases different data were used for each study. The mean and maximum interstudy differences for f_u_ were 3.04- and 15-fold, respectively. Also in this case, replacement of selected f_u_ and C_bl_/C_pl_ with average values resulted in large decrease of R^2^ and increase of prediction errors.

Another major pharmaceutical company reported 5.9- and 19-fold mean and maximum differences in hepatocyte CL_int_-estimates between studies (n=19). In 91 % of cases, different f_u_- and C_bl_/C_pl_-data were used for each study. The mean and maximum interstudy differences for f_u_ were 1.95- and 14.5-fold, respectively. Corresponding estimates for C_bl_/C_pl_ were 1.16- and 1.63-fold, respectively.

Thus, it is common that researchers and companies select different reference estimates for different studies. An apparent consequence is that exaggerated predictive performances are demonstrated. With the use of mean values from different sources/studies a more accurate and less biased picture is reached and shown.

Another way to improve the R^2^-values is to exclude/remove non-quantifiable compounds (e.g. CL_int_ and f_u_<LOQ) and compounds with high lipophilicity (low solubility and poor dissolution) and low recovery. Such compounds are seldom included and reported. Atenolol is a compound with negligible *in vivo* metabolism, but with reported moderate/high *in vitro* CL_int_. As shown above, inclusion of results for this compound has a major negative effect on the R^2^ for plasma CL_H_ (3-fold higher R^2^ when excluding atenolol). There are also examples of studies where compounds with low f_u_ are avoided, which reduces uncertainty and errors for CL-predictions.

The unbound fraction in hepatocyte incubations (f_u,inc_) is another source of variability and uncertainty. The f_u,inc_ for the 32 selected compounds averages 0.80. Only, two of them has a f_u,inc_ < 0.5 (diazepam; f_u,inc_ = 0.15, montelukast; f_u,inc_ = 0.0002). The mean and maximum of ratios between highest and lowest reported estimates were 1.47- and 2.73-fold, respectively. This was more and less extensive than for C_bl_/C_pl_ and f_u_, respectively.

Based on the average choice of f_u_, C_bl_/C_pl_ and f_u,inc_ data a 4-fold difference (2.29 • 1.15 • 1.47) of predicted CL_H_ can be produced. The corresponding maximum difference is 65-fold (15 • 1.59 • 2.73).

### Comparing predictions based on *in vitro, in silico* and animal *in vivo* data

For the 29 compounds for which *in vitro* data-based F-predictions resulted in a R^2^ of 0.20 (slope=0.57; intercept=0.23; see above) *in silico* predictions (using methodology by Prosilico AB) reached a Q^2^ (R^2^ for forward-looking predictions; compounds not included in training sets of models) of 0.58 (slope=0.82; intercept=0.13; Figure 8). Thus, the *in silico* methodology clearly outperformed the *in vitro*-to-*in vivo*-methodology with average data from many sources.

**Figure 8.**
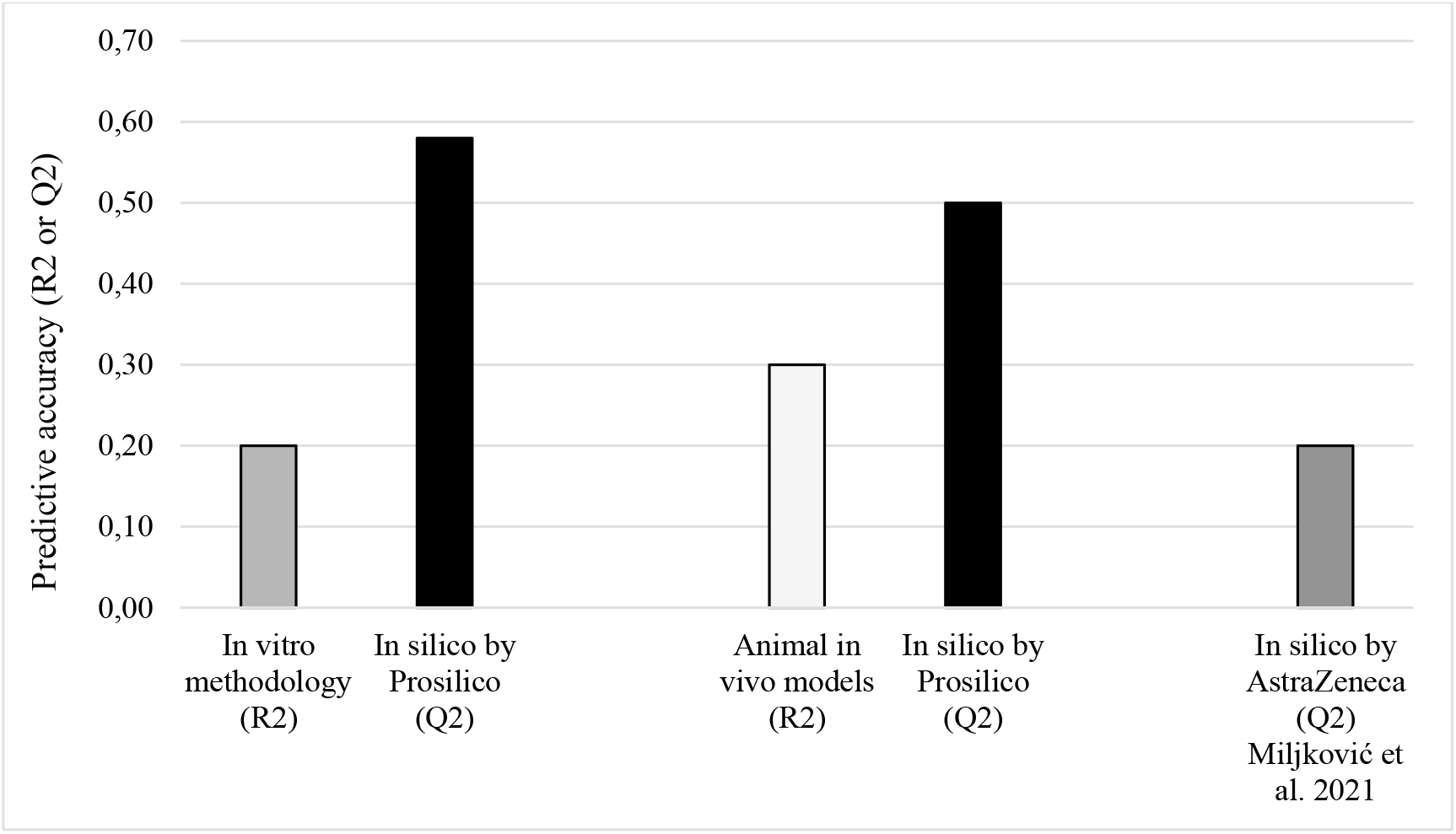
Predictive performance of *in vitro* (R^2^), *in silico* (Q^2^) and animal *in vivo* models (R^2^) for F. Head-to-head comparisons were done for *in vitro vs in silico* (n=29) and animal *in vivo vs in silico* (n=156).

In a previous study, this *in silico* method outperformed animal models (mice, rats and dogs) in a head-to-head comparison (Q^2^=0.50 for *in silico* method *vs* R^2^=0.30 for animal models combined; n=156) (Fagerholm et al. 2021b) (Figure 8). The *in silico* method also produces a higher Q^2^ compared to a new *in silico* method by AstraZeneca (Q^2^ ca 0.1-0.3) (Miljković et al. 2021) (Figure 8).

The poorer predictive performance of *in vitro* methodology *vs* animal models is in line with previous findings (Poulin et al. 2011).

Other advantages with the *in silico* method is that it is devoid of prediction data selection bias and can predict CL_H_ and F for compounds out of reach for *in vitro* methods, such as those with CL_int_ and f_u_<LOQ and high lipophilicity, low solubility, poor dissolution and low recovery, and for low permeability compounds.

## Conclusion

The combination of data from many laboratories and the use of mean values resulted in reduced selection bias and predictive accuracy. Overall, the predictive accuracy (here R^2^) for log CL_int_, log CL_H_ and F was low to moderately low (0.08-0.32). The halved R^2^ compared to individual studies where high performance was demonstrated seems to be explained be selection bias (enabled by large data variability). Animal *in vivo* models, and in particular, *in silico* methodology, outperformed *in vitro* methodology for the prediction of F in man. The encouraging results for *in silico* methodology is a major step forward in the process of replacing an reducing animal experiments in pharmacokinetics and drug discovery and development.

